# Cognitive control networks in human and macaque

**DOI:** 10.64898/2026.02.24.707621

**Authors:** Valentina Mione, Freja Holm Prins, Moataz Assem, Urs Schüffelgen, Søren Kyllingsbæk, Mark J. Buckley, Daniel J. Mitchell, John Duncan

## Abstract

A much-replicated finding in human brain imaging is a distributed “multiple-demand” or MD system, increasing in activity for many kinds of cognitive demand, and centrally involved in cognitive control. MD regions are proposed to encode a distributed mental model of critical task events, bound together in the roles and relationships needed to direct action selection. Though previous data hint at a corresponding network in the macaque, there has been no direct comparison to human data. Here we used functional magnetic resonance imaging to measure whole brain activation in a multi-step saccadic maze task, compared to a control requiring similar moves but without goal-based decisions. Human data were a close match to the canonical MD network, extended to include adjacent regions and in particular much of the canonical dorsal attention network. Data from two monkeys suggested correspondences in dorsomedial frontal, insula/orbitofrontal and posterior temporal cortex. In lateral frontal cortex there was just a single, largely dorsal activation patch, in contrast to multiple distinct human patches. In one animal, as in human, there was accompanying activation of dorsal and medial parietal cortex. In macaque as in human, together with previous data, our findings suggest an extended and strongly interconnected brain network recruited by increased cognitive challenge.

## Introduction

Among the most stable findings of human brain imaging is a “multiple-demand” or MD system, defined by increased activation for a very wide range of cognitive demands (Duncan and Owen, 2000) (Duncan, 2010) (Duncan et al., 2020). With its broad recruitment, the MD system is thought to play a central role in attentional or cognitive control (Cole and Schneider, 2007) (Duncan et al., 2020). The high-resolution methods of the Human Connectome Project (HCP) show nine distinct MD patches (Assem et al., 2020) (Assem et al., 2024), distributed across lateral and dorsomedial frontal cortex, anterior insula, dorsal and medial parietal cortex, and posterior temporal cortex (Figure 1a). Within these patches, based on the cortical parcellation of (Glasser et al., 2016), Assem et al. (Assem et al., 2020) defined a set of ten “core” MD parcels (Figure 1b), identified by the strongest converging response to multiple cognitive demands, and strong interconnections indicated by resting state correlations. Common activity for different demands, however, usually extends beyond these core MD parcels, in particular to adjacent regions of dorsal attention (DA) and cingulo-opercular (CO) networks (Ji et al., 2019) (Assem et al., 2024) (Figure 1b). For different demands, the precise activation pattern shifts somewhat within the same nine MD patches, often with differential recruitment of individual DA and CO parcels (Assem et al., 2024), but with strong overlap between demands (Figures 1c & d).

**Figure 1.**
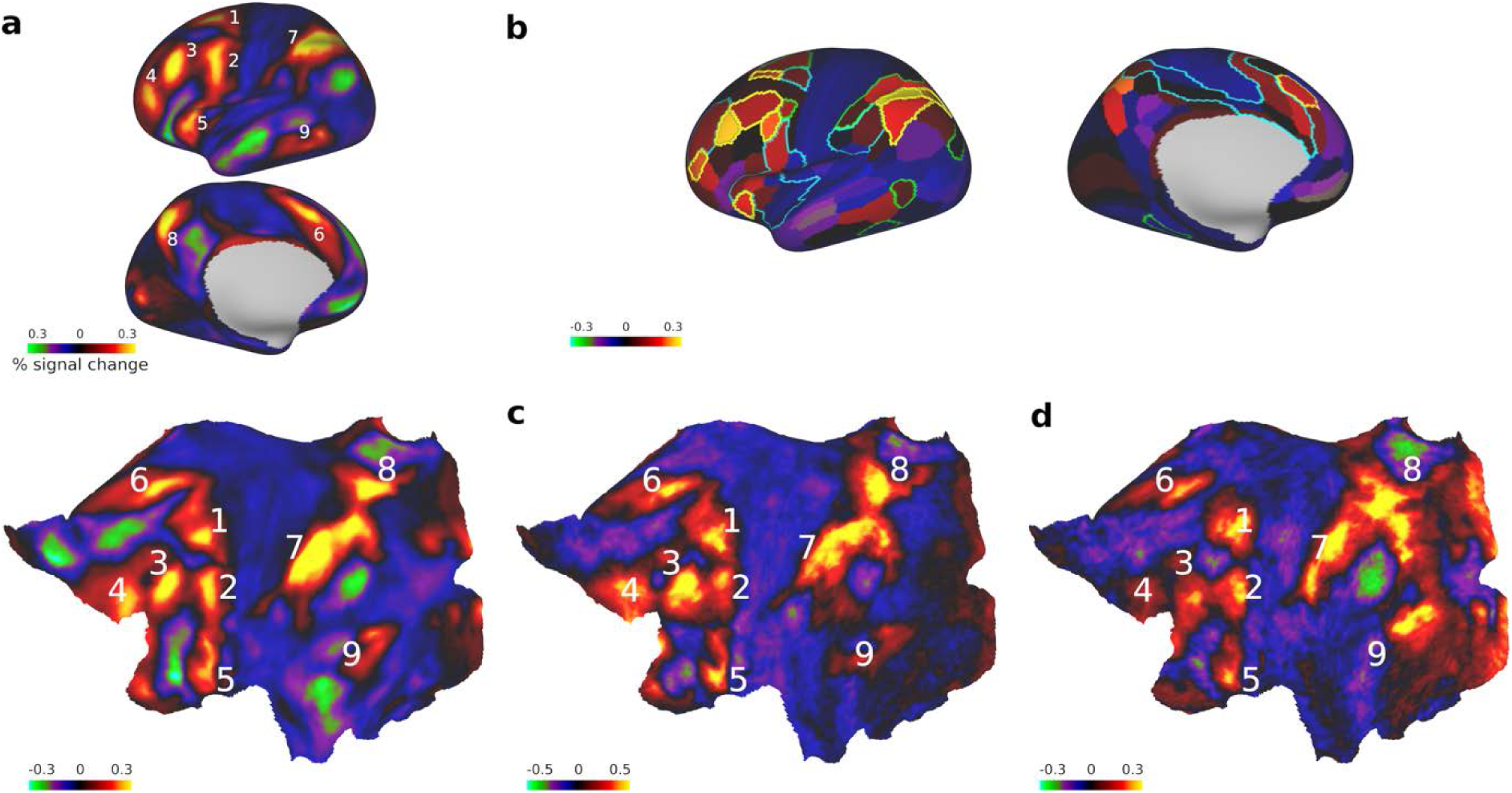
Human multiple-demand (MD) system. (a) 9-patch MD system based on mean activation across three dissimilar demands. Data from Human Connectome Project (Assem et al., 2020). Top – inflated cortical surface; bottom – cortical flatmap. (b) Same data – average activation in individual cortical parcels from (Glasser et al., 2016). White outlines – core MD parcels (Assem et al., 2020); green outlines – dorsal attention network (Ji et al., 2019); blue outlines – cinguloopercular network (Ji et al., 2019). (c) and (d) 9-patch MD system in independent data – (c) n-back working memory (Assem et al., 2024); (d) mental rotation (Faber et al., 2024).

Cognitive control must rest on an internal model of a task or situation, with parts combined in the correct roles and relationships (Kimberg and Farah, 1993) (Duncan, 2025). It is the contents of this task model that indicate correct behavior for a current goal. Studies using multivoxel pattern analysis show that, across the human MD network, there is extensive coding of task-relevant information, including stimuli, rules and responses (Woolgar et al., 2016). Elsewhere, we have suggested that the widely distributed parts of the MD network give access to many different kinds of information, while strong interconnectivity allows different kinds of information to be integrated into a task model (Duncan et al., 2020).

A similar distributed network likely exists in the macaque. Electrophysiological studies, certainly, show selective neural responses for many kinds of task event in lateral frontal, dorsomedial frontal and parietal cortex (Amemori et al., 2015) (Cavanagh et al., 2018) (Chafee and Goldman-Rakic, 1998) (Goodwin et al., 2012) (Johnston et al., 2007) (Kadohisa et al., 2020) (Loonis et al., 2017) (Miller and Cohen, 2001) (Salazar et al., 2012). Provisional support comes from a handful of imaging studies. One comparison between pro- and antisaccades showed increased activity in frontal eye field, anterior cingulate and intraparietal sulcus for the more demanding antisaccade (Ford et al., 2005) (Ford et al., 2009). A study of task switching suggested activation of lateral frontal cortex, anterior cingulate, and a region of orbitofrontal cortex close to anterior insula (Premereur et al., 2018). Measures based on functional (Mitchell et al., 2016) and structural (Karadachka et al., 2023) connectivity have also been used to estimate putative components of a macaque MD system, including regions of lateral frontal, dorsomedial frontal, inferior parietal and medial parietal cortex. In the absence of a direct comparison between species, however, it remains uncertain whether the macaque brain contains a functional analogue to the distributed human MD system.

To address this uncertainty, we collected fMRI data from humans and macaques performing a similar on-screen maze task. In the human, using HCP methods, we show the usual nine-patch pattern of brain activity associated with increased cognitive demand. In the macaque, we also observe widespread demand-related brain activity, and ask how well it matches the human nine-patch pattern.

## Results

### Human

The human maze task is illustrated in Figure 2a. With a series of saccades, the task was to navigate from a central start location (Figure 2a, white circle) to one of four possible goal locations (Figure 2a, blue circles). Along the route, the current location was defined as the current fixation point; permitted fixation locations were the intersections and corners of maze paths, and each saccade moved just one step through the maze, i.e. from the current location to one of the immediately adjacent locations. An initial display indicated the goal for the current problem (Figure 2b, top), always one of the four shown in Figure 2a. At each step towards this goal, a choice display indicated possible moves (Figure 2b). Of the locations immediately adjacent to the current location, two were colored yellow, indicating that they were open (available for selection), while the remainder were colored red, indicating that they were blocked. A delay of 0.4 or 1.0 s preceded a go signal (Figure 2b, blue dot in white circle), indicating that the next saccade could be made. The correct choice was then one of the two open (yellow) locations, if possible moving closer to the goal. This procedure was repeated for a series of steps, until the goal was finally reached and a reward signal was received. Depending on the available choices at each step, the goal could be reached directly, in just two steps, or less directly, in up to six steps including forced steps away from the goal.

**Figure 2.**
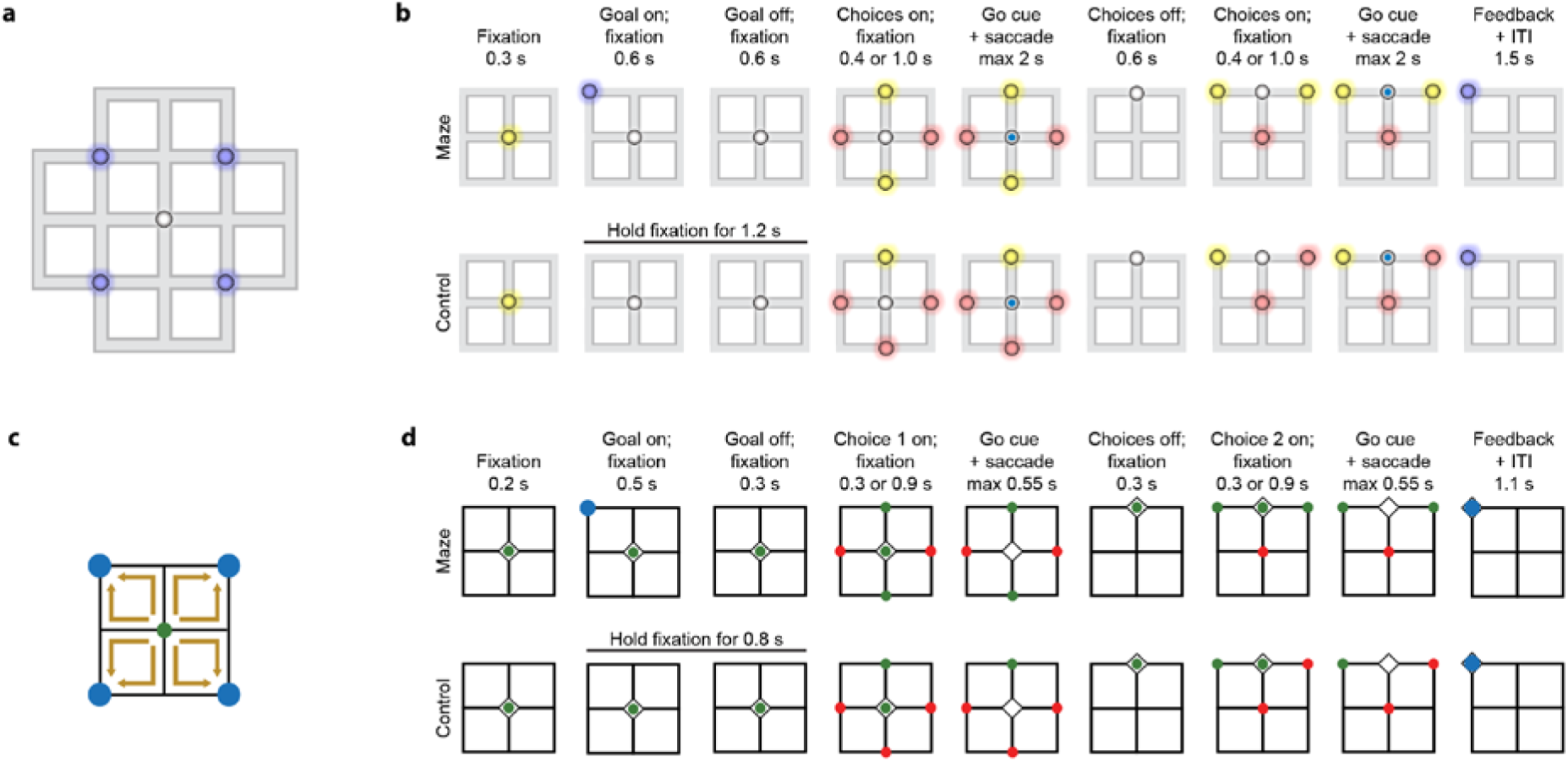
(a) Human maze. White circle – initial fixation location; blue circles - possible goal locations. (b) Human maze stimuli and timings. Upper: Example problem in the maze condition (outer parts of maze not shown, though present on screen). On each step, yellow dots marked open locations available for selection, with red locations blocked. On receipt of the go signal (blue dot inside white circle), participants made a saccade to one of these yellow locations. The task was to choose the shortest route to the goal. In this example, the goal is reached directly in two steps. In other problems, longer routes were created by enforcing some moves that did not move closer to the goal. Timings for each step were identical. Lower: Control condition. (c) Monkey maze. All problems required just two steps to the goal (brown arrows). (d) Monkey maze stimuli and timings (upper – maze; lower - control). Green locations – open; red locations – blocked.

In a control task (Figure 2b, bottom), there was no goal and only a single location was open (yellow) at each step. Again, the participant made a series of steps through the grid, but in this case, simply following the route prescribed, until a reward signal was given when reaching one of the (unmarked) goal locations after between 2 and 6 steps.

Behavioral data are shown in Figure 3a. Responses were recorded when the eye reached one of the open or blocked locations, with saccades elsewhere ignored. For each step through the maze, accordingly, there were five possible outcomes (correct, too fast, too slow, move to blocked location, move to wrong open location). Data in Figure 3a are just for steps in which the two open locations included one that was correct (towards the goal), one that was not (away). In both maze and control tasks, the majority of errors were either too fast or slow, with few wrong choices among the indicated open and blocked locations.

**Figure 3.**
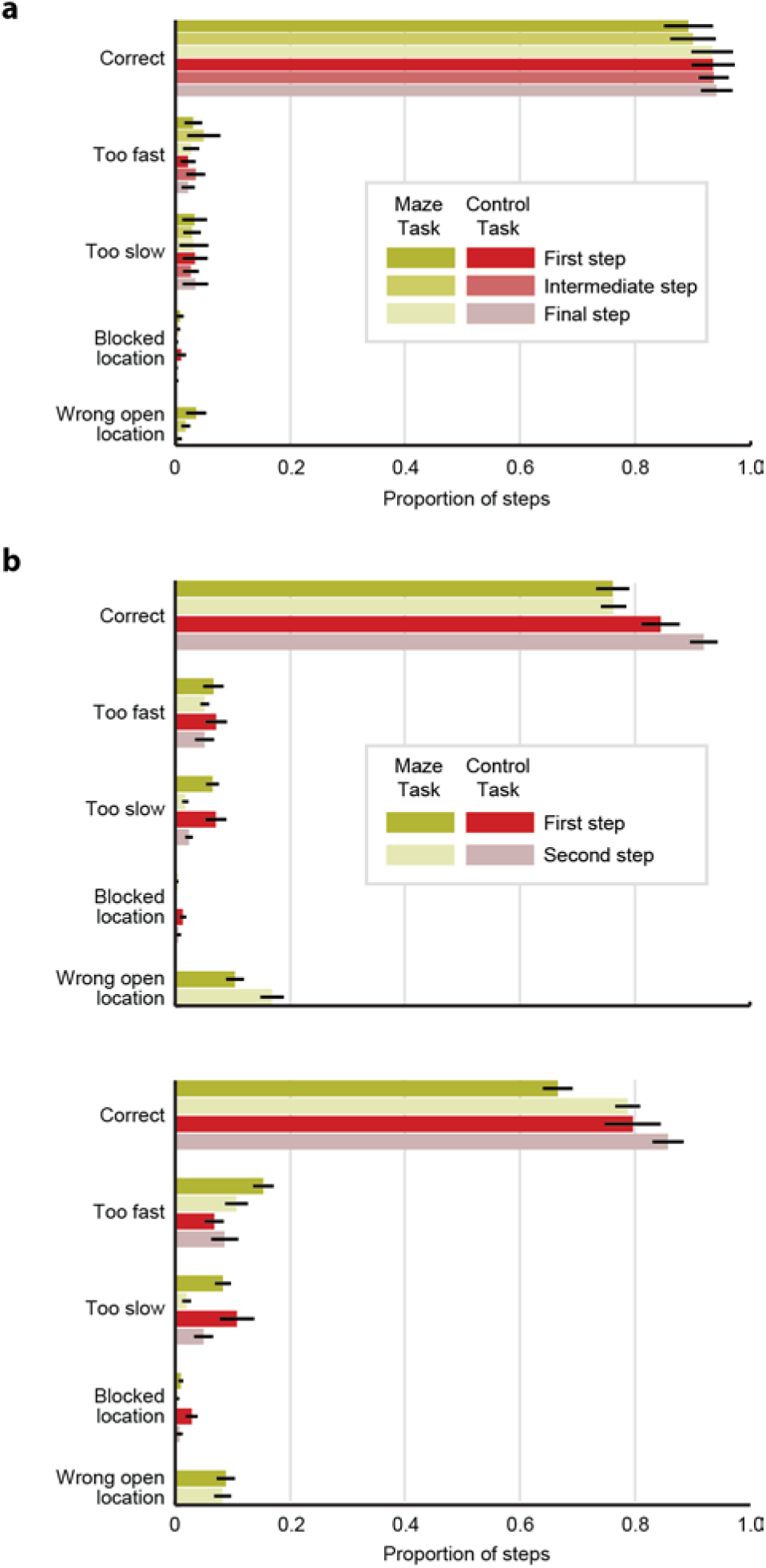
Behavioral data. (a) Human. Bars show 95% confidence intervals over participants. (b) Monkeys A (top) and B (bottom). Bars show 95% confidence intervals over sessions.

Functional magnetic resonance imaging (fMRI) data were collected in blocks of problems of a given type (maze or control). Each block lasted for 24 s, plus the time required to complete the final problem. Every five consecutive blocks contained two blocks each of the maze and control task, along with one additional block (not analysed) of a simple central fixation task.

Whole brain data for the contrast of maze vs. control are shown in Figure 4. Figure 4a shows whole-brain activation, on semi-inflated cortical surfaces for each hemisphere (center), with cortical flatmaps to either side. Figure 4b shows average data for the two hemispheres, related to the cortical parcellation of (Glasser et al., 2016). Colors for each parcel indicate mean activation across all vertices within that parcel. Figure 4c shows mean activations for each of 12 cortical networks, defined by (Ji et al., 2019) using the (Glasser et al., 2016) parcellation. At the left is mean activation for the 10 core MD parcels defined by (Assem et al., 2020), all of which belong to the frontoparietal network (FPN) from (Ji et al., 2019) (Figure 1c, yellow). Results for remaining FPN parcels are shown separately. In Figure 4b, colored outlines mark parcels of core MD (yellow), dorsal attention (DA, green) and posterior multimodal networks (PM, lilac).

**Figure 4.**
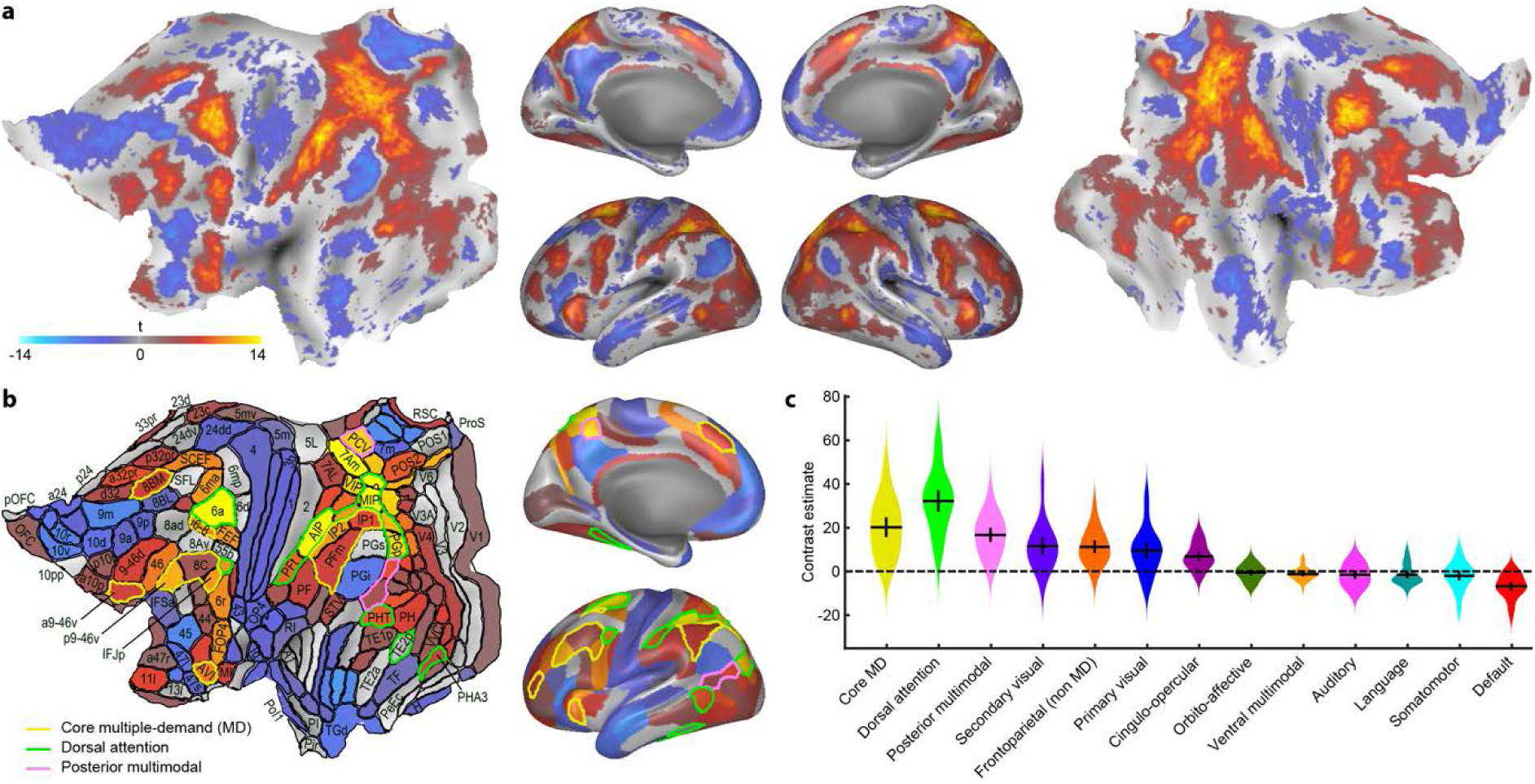
Contrast of maze minus control: Human data. (a) t values (thresholded at FDR<0.05) projected onto partially inflated cortical surface (center) and cortical flatmaps (sides). (b) Mean t value (for regions with FDR < 0.05) in individual cortical parcels from (Glasser et al., 2016). Yellow outlines – core MD parcels (Assem et al., 2020); green outlines – dorsal attention network (Ji et al., 2019); pink outlines – posterior multimodal network (Ji et al., 2019). Parcel labels from (Glasser et al., 2016); some omitted to avoid clutter. (c) Mean values in the 12 cortical networks defined by (Ji et al., 2019). Data are shown separately for core MD and other parcels from the frontoparietal network defined by (Ji et al., 2019). Horizontal and vertical black lines show mean and 95% confidence intervals; violins show distribution across participants.

In broad outline, activations for this maze vs. control contrast strongly resembled the canonical MD system (compare Figure 4a to Figure 1). The same six patches were observed in lateral frontal cortex, anterior insula and dorsomedial frontal cortex. Maze activations also encompassed the two parietal MD patches, though with substantial additional spread on both lateral and medial surfaces. A posterior temporal activation was also a close match to the posterior temporal patch from (Assem et al., 2020).

Parcel-wise results in Figure 4b confirm positive activations in all core MD parcels, substantially extending into adjacent regions. Strong activations in particular, were seen in many parcels of the DA network, with the next strongest average activity in PM (Figure 4c). Additional activations extended across a large region of medial parietal cortex, a number of dorsomedial frontal parcels adjacent to core area 8BM, and lateral frontal regions immediately dorsal and postero-ventral to core MD regions. Strong activation was also seen in the frontal eye field (Figure 4b, FEF).

For a closer match to the macaque task (see below), the analysis was repeated using data just from two-step problems. The results (Supplementary Figure 1) were closely similar to results from the main analysis, with the expected reduced power. For comparison, we also examined data from a version of the task with manual rather than saccadic responses (see Methods). Again, results were closely similar (Supplementary Figure 2).

### Macaque

The monkey task was a simplified version of the human maze (Figure 2c). The goal was always reached in two steps, the first moving to one of the four locations immediately adjacent to the central start location, and the second reaching the goal. Again, animals worked continuously through problems for the duration of each block, which lasted a minimum of 24 s, plus the time required to complete the final problem.

Behavioral data are shown in Figure 3b. Both animals had lower accuracy than humans, but an otherwise similar distribution of errors. Again, in particular, there were few choices of blocked locations, and many more correct than incorrect choices between the two open locations, indicating strong control of behavior by the remembered goal.

For imaging analysis, because timing errors often indicated poor task engagement, we took data just from problems that ended with saccade to an open or blocked location, made in the correct time window. For the main analysis, to increase power, data from the two animals were combined (see Methods), and data from the two hemispheres were averaged.

In Figure 5, the contrast of maze vs. control is shown on flatmap (left) and semi-inflated surfaces (right) of the left hemisphere. Parcellation is according to the atlas of (Saleem et al., 2021), projected onto the F99 surface (Van Essen et al., 2012).

**Figure 5.**
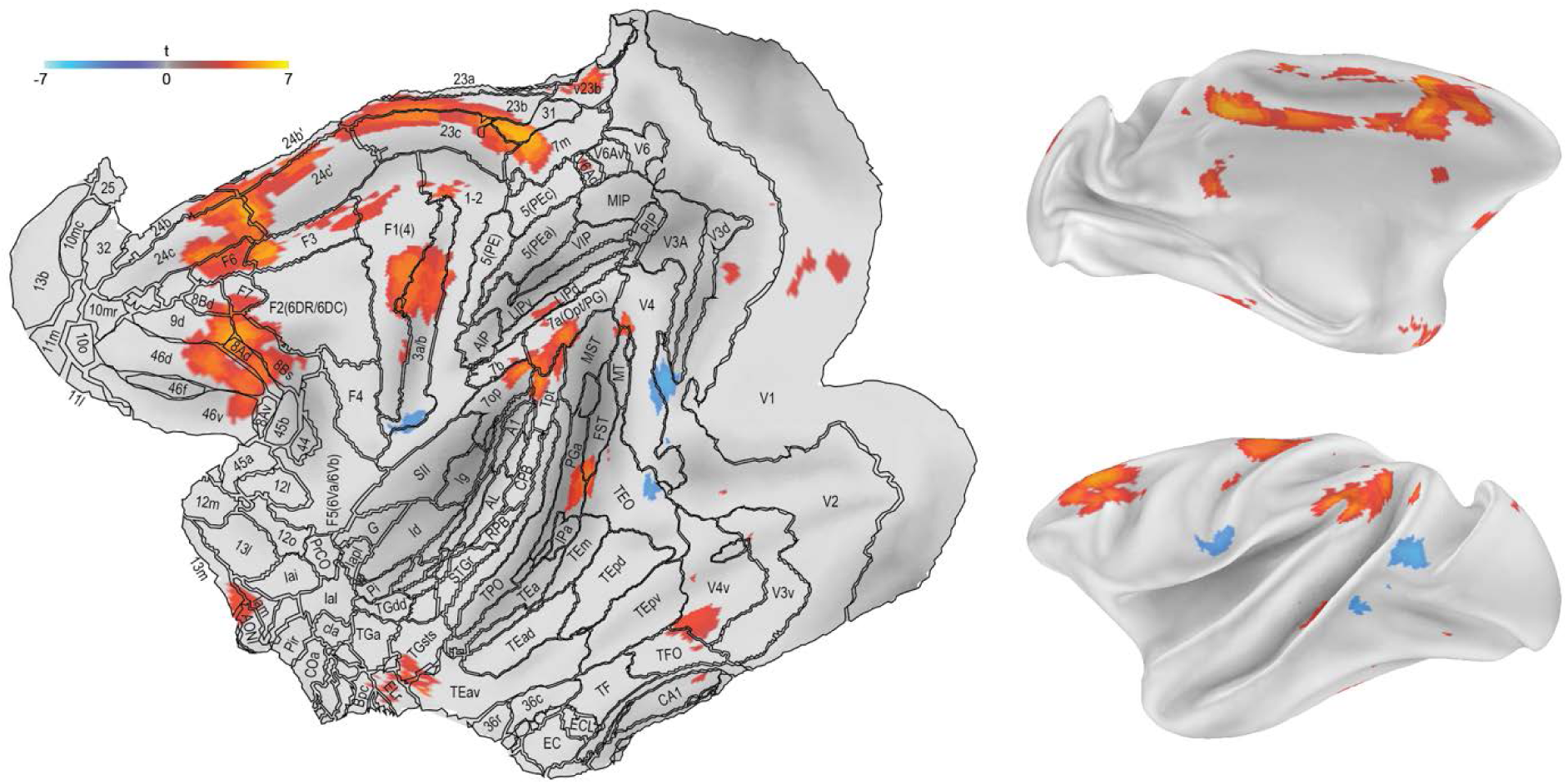
Contrast of maze minus control: Monkey data. Left – cortical flatmap, with projected t values (thresholded at FDR < 0.05). Parcel labels according to (Saleem et al., 2021); some omitted to avoid clutter. Right – partially inflated cortical surface.

In several respects, there was good correspondence between monkey and human data. On the dorsomedial frontal surface, there was a large patch of activation centered on areas F6, 24c and 24c’, corresponding to the monkey’s preSMA and anterior cingulate. On the lateral parietal surface, a patch of activation extended over parts of areas 7a (Opt/PG) and 7op, extending into adjacent Tpt. In agreement with the human data, there was also a patch of activation in medial parietal cortex, extending from 7m to adjacent parts of 23b/c and 31. In the insula, there was activation in Iam, extending into adjacent regions of orbitofrontal cortex. In occipitotemporal cortex there was a patch of activation in anterior V4v, tentatively corresponding to the posterior temporal patch in human data.

Correspondence was less clear on the lateral surface of the frontal lobe. In the monkey, contrasting with the multiple, clearly separable, activation patches in the human, there was just one large posterior dorsal patch extending from F7 (dorsal premotor) through 8ad (frontal eye field) to dorsolateral frontal cortex (9d, 46d) dorsal to the principal sulcus, and within its dorsal bank. There was modest extension into the ventral bank of the sulcus (46v), but not to the ventrolateral frontal surface. Differences between human and macaque were also notable in dorsal parietal cortex, with strong activation of DA regions in the human, not visible in the macaque.

Finally, there were further activation patches in the monkey not matched in human data. Most conspicuously, in the monkey, there was a large activation patch in sensorimotor cortex (3a/b, extending into ventral F1), and additional scattered activations close to the temporal pole.

In a control analysis, we included data just from correctly completed problems. Results (Supplementary Figure 3) closely resembled those from the main analysis. In a second analysis, we reduced the smoothing kernel from 4 mm full width half maximum (FWHM) (see Methods) to 3 mm, both commonly used in monkey fMRI studies (e.g. (Gandaux et al., 2025; Miyamoto et al., 2017)), and approximately matching the smoothing required to match electrophysiological to fMRI maps (Issa et al., 2013). Though activations were more restricted at 3 mm smoothing, their cortical distribution was similar (Supplementary Figure 4).

Finer breakdowns of the data are shown in Supplementary Figures 5 and 6. Activations were closely similar for left and right hemispheres (Supplementary Figure 5). Separate data from the two animals, however, showed a number of differences (Supplementary Figure 6a). Most strikingly, except in lateral frontal cortex, activations were stronger and more extensive for one animal. Ony this animal showed parietal lobe activations. Of note, in one of the two animals, there was extensive dorsal parietal activation, not visible in the combined data; while in the other, lateral frontal activation spread ventrally, to include regions 45b and 44. Even at a permissive threshold of t > 1.5, a conjunction analysis (Supplementary Figure 6b) showed no common activation in parietal cortex.

**Figure 6.**
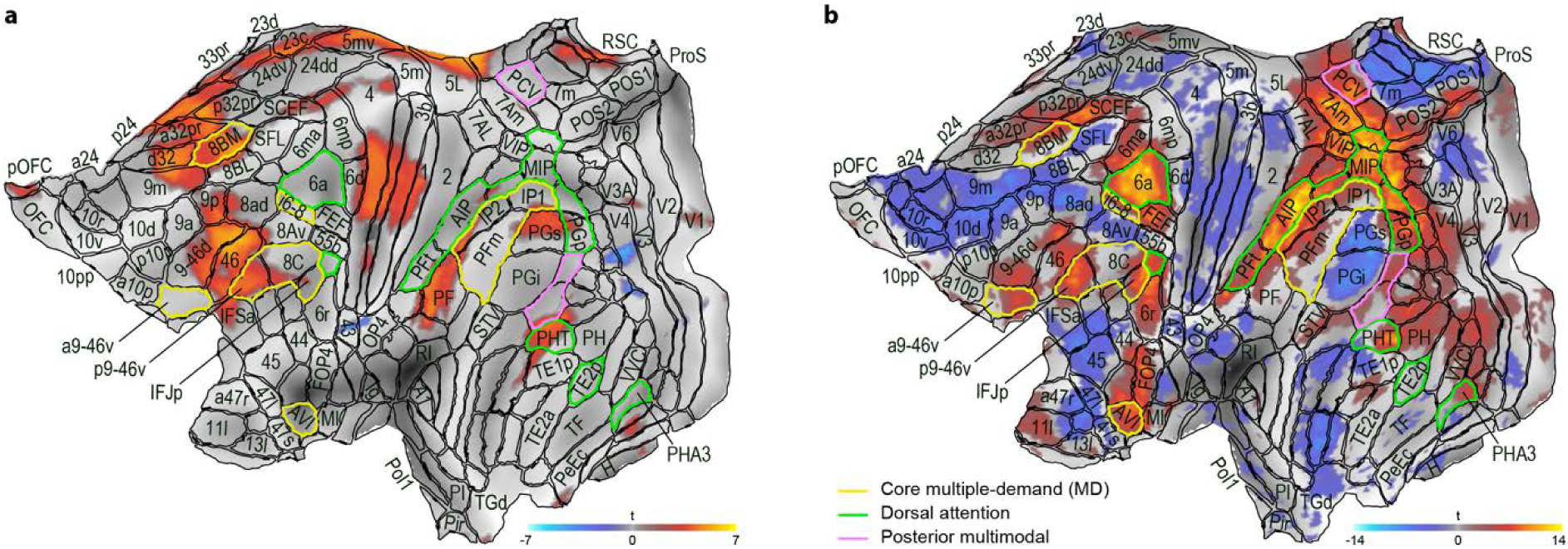
(a) Monkey data: Contrast of maze minus control warped to human cortical surface. (b) Human data for comparison.

A variety of methods have been proposed for approximate warping of the macaque to the human brain. Figure 6a shows macaque activations from Figure 5 warped by a method based on resting state fMRI data (Xu et al., 2020), with human activations repeated for comparison (Figure 6b). With this approximate warping, lateral frontal activations show substantial overlap between species. As noted in the main analysis, dorsal parietal activation is absent in the macaque data. In this warped version, insula/orbitofrontal activation in the macaque is placed in human pOFC (top left of flatmap), perhaps reflecting limitations of the warping method in this part of the brain.

### Subcortical activation

In the human, cortical MD activity is generally accompanied by focal activations in caudate, thalamus and cerebellum.(Assem et al., 2020) The same was true in the present human data (Figure 7a), and in the monkey, regions of activation were also seen in cerebellum and caudate (Figure 7b).

**Figure 7.**
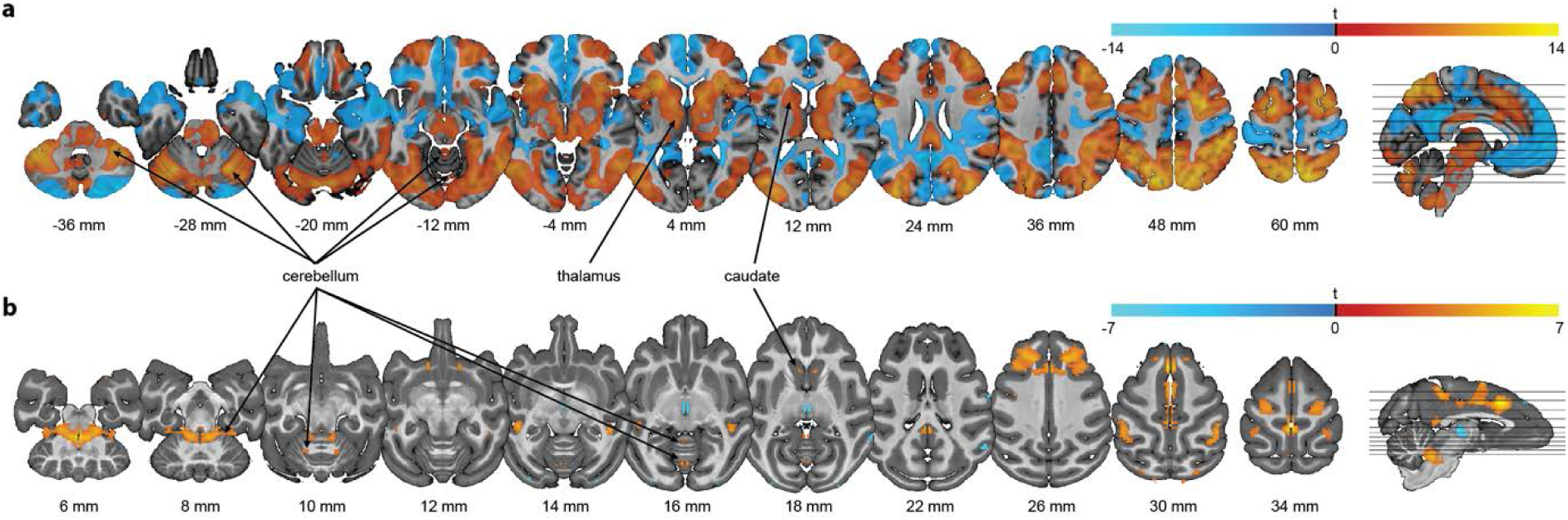
Contrast of maze minus control shown on horizontal slices (slice locations shown on right). (a) Human. (b) Monkey.. Threshold p < .05, FDR corrected over the whole brain.

## Discussion

Though precise correspondences between brain networks in human and monkey are uncertain, the whole-brain data of fMRI can provide a useful comparison. Here, we used fMRI in human and monkey to compare brain activation during a multi-step spatial maze task.

In humans, as expected, we found a highly specific, distributed pattern of brain activity, incorporating the canonical MD system, and extending to adjacent regions of DA and other networks. MD activations strongly resembled those broadly associated with increasing cognitive demand, while DA and visual activations were likely associated with the specific visuospatial content of the maze task.

In several respects, monkey data were a good match to human MD activations. Patches of activation were seen in dorsomedial frontal cortex, insula and adjacent orbitofrontal cortex, and a small patch in occipitotemporal cortex. Conspicuous activation was also seen in lateral frontal cortex, though in contrast to several distinct patches in the human, monkey activations were seen only in one large region extending from premotor cortex to the banks of the principal sulcus. In the human, the contrast of maze versus control showed strong activation of parietal cortex, including regions of both MD and DA networks. In the monkey, strong parietal activation was seen in only one of the two animals (Supplementary Figure 6a).

In human imaging, the MD system is defined by overlapping activation for varied cognitive demands. In the present work, in contrast, we used just a single task. There is some overlap between our putative monkey MD network and previous findings, including activations of intraparietal sulcus and anterior cingulate reported for antisaccades compared to prosaccades (Ford et al., 2005) (Ford et al., 2009), and of lateral frontal cortex, anterior cingulate and orbitofrontal cortex close to insula for task switching compared to repeat (Premereur et al., 2018). Though a wider variety of tasks would be desirable, these overlaps suggest that, in the macaque brain as in the human, an extended MD network is recruited by diverse cognitive challenges.

Our results may be compared to previous estimates of a macaque MD system, based on either functional or structural connectivity. In the human brain, MD regions show strong functional connectivity, indicated by correlated activity time-courses in resting state data (Assem et al., 2020). Previously, we searched for a similar network based on functional connectivity data from anesthetised monkeys (Mitchell et al., 2016). On the one hand, strong connectivity between regions of lateral frontal, dorsomedial frontal and inferior parietal cortex was coarsely in line with current findings. Often these strongly-connected regions were close to or overlapping with activated regions in the maze task. At the same time, there were notable differences between networks defined by these two methods. The functional connectivity network, for example, included a larger region of premotor cortex, extending back into primary motor cortex, and ventrally well beyond the principal sulcus. This network, furthermore, included neither orbitofrontal/anterior insular nor medial parietal cortex. Plausibly, a network underlying high-level cognitive control is only partially visible in time course correlations in anesthetised animals. A second study identified macaque regions that most strongly resemble human MD regions in terms of structural connectivity to major white matter tracts (Karadachka et al., 2023). Again there was coarse overlap with the current network, including regions in lateral frontal, dorsomedial frontal, inferior parietal and medial parietal cortex, but again, this coarse agreement was accompanied by notable differences. Once more, for example, the structural connectivity network included a much larger region of lateral frontal cortex. Plausibly, connectivity to major white matter tracts gives only a partial clue to functional specialisation, especially for regions in close anatomical proximity.

In the present macaque data, data combined across the two animals showed maze-related lateral frontal activation that was largely restricted to dorsal regions, with minimal extension onto the inferior convexity (ventrolateral frontal cortex, vlPFC). Extension into vlPFC was seen only in a single animal (Supplementary Figure 6a). Others have suggested that vlPFC plays a core role in cognitive and attentional control (Passingham and Wise, 2012) (Trambaiolli et al., 2022), and in a recent electrophysiological study, we found that coding of both spatial and nonspatial task information was stronger in ventral than dorsal frontal cortex (Kadohisa et al., 2023). In a spatial maze task too, electrophysiological recordings show a strong role for vlPFC in rapid discrimination of relevant sensory input, including initial goal and subsequent choice stimuli (Mione et al., 2025). In anesthetised macaques, the functional connectivity profile of vlPFC regions also shows resemblance to the proposed MD network, including connectivity in dorsomedial frontal and lateral parietal cortex (Neubert et al., 2014) (Mitchell et al., 2016).

The monkey data also showed regions not mirrored in the human. Most conspicuous was a region of sensorimotor cortex. The imaging situation, of course, may often be different for monkey and human. While humans lie still, monkeys do not, and possibly, the amount of movement depends on task demands. Other differences in procedure may have limited similar brain activity in the two species. While humans learned and performed their task in a single session, macaques were trained in a series of steps over an extended period (see Methods). In macaque sessions there were only two-step problems, while in humans, even two-step problems may have been influenced by a surrounding context of more complex problems. Evidently, similarities of human and macaque activations survived these large procedural differences.

Especially given differences in results for the two animals, questions remain concerning possible species differences. As noted above, striking components of the human MD network (vlPFC and parietal cortex) were each seen in only one of the two animals. Limited statistical power, limited difficulty of the monkey task and long training may all have contributed to negative results for single animals. Additional data, preferably involving a wider range of task demands, would be needed to address such limitations. At the same time, results from single animals tell against the possibility of genuine species differences in these regions.

The existence of MD regions, recruited as a network to address diverse cognitive challenges, does not imply homogeneity of function, either between regions or within a single region. Between tasks, human data show partial functional specialisation of different MD regions (Assem et al., 2020). Within a single patch, there are gradients of activation for any given task demand, with the magnitude and direction of these gradients varying from one task to another (Assem et al., 2024). In the monkey, too, functional connectivity profiles for different regions of lateral frontal cortex show strong family resemblances, but also quantitative variations (Neubert et al., 2014). Differing anatomical connections likely give MD regions differential access to information of different kinds (Passingham and Wise, 2012); at the same time, strong interconnectivity may bring substantial information sharing and co-recruitment.

In putative MD regions defined here, electrophysiological studies show coding of many different kinds of task-relevant information. This is amply documented in lateral frontal, dorsomedial frontal, insula/orbitofrontal, premotor and lateral parietal cortex(Amemori et al., 2015) (Cavanagh et al., 2018) (Chafee and Goldman-Rakic, 1998) (Goodwin et al., 2012) (Johnston et al., 2007) (Kadohisa et al., 2020) (Loonis et al., 2017) (Miller and Cohen, 2001) (Mizuhiki et al., 2007) (Nakajima et al., 2022) (Salazar et al., 2012) (Vallentin et al., 2012) (Yang et al., 2022). These results mirror human fMRI data, showing extensive MD encoding of a task’s stimuli, rules and responses (Woolgar et al., 2016). In monkey as in human, we suggest that a distributed MD network assembles information of many kinds into a distributed task model, needed to represent task events and to choose appropriate behavior.

## Methods

### Human: Participants

39 participants took part, of whom two were not analysed owing to missing or low quality structural scans, and three were excluded owing to poor performance (see below). The 34 participants (22 female) in the analysed sample had a mean age of 30.2 years, with normal or corrected-to-normal vision, and with no history of neurological or psychiatric disorders. Participants provided informed consent and were paid for their time. The experiment was conducted in accordance with ethical approval from the Cambridge Psychology Research Ethics Committee.

### Human: Maze task

The experimental task was programmed in Matlab (The Mathworks Inc.) using Psychtoolbox version 3.0.14 (Kleiner et al., 2007). In two of four runs, the subject of the present report, participants responded by making saccades to a sequence of target locations, while their gaze was monitored using an eye tracker (EyeLink 1000; SR Research Ltd). In the other two runs (not analysed here, except for comparison data in Supplementary Figure 2), tasks were identical except that responses were made by button press. Saccadic runs were either the first two or the last two, alternating across participants. In two participants, one of the saccadic runs was missing due to technical issues. When responding using saccades, a location was deemed fixated when gaze had remained within 1.6 degrees of visual angle (dva) of the specified coordinate for 0.3 s.

The experiment used a blocked design, with each run containing 25 blocks, in five groups of five. Each group contained two blocks of the maze task, two of control, and one (not analysed) of fixation. The fixation block was always the third in the group, with one block each of maze and control before the fixation block, one block each following it, and the order of maze and control blocks counterbalanced across runs. Within each block, problems were presented one after another for 24 s, after which the current problem was allowed to finish before proceeding to the next block. Blocks comprised 1-6 problems (median 3), depending on the number of required saccades and the performance of the participant. Each run lasted approximately twelve minutes.

For every problem, participants were presented with a grey grid, whose width spanned approximately 11.5 dva, on a white background (Figure 2a). At the start of each problem, a circular yellow fixation dot (0.7 dva) was positioned at the center of the grid (Figure 2b). The participant was required to fixate this dot for 0.3 s, whereupon it turned white, whilst fixation was maintained for a further 1.2 s.

During this period, events diverged for maze and control tasks. In the maze task, for 0.6 s, one of the four possible goal locations was highlighted by a blue circle (Figure 2b), then disappeared for a further 0.6 s. The goal location was always one of the four shown in Figure 2a. At the end of this 1.2 s period, two movement options appeared as yellow circles at grid intersections immediately adjacent to the fixation location; the other adjacent locations were marked by red circles, indicating them as blocked. The participant was to maintain fixation for a further 0.4 or 1.0 s (equal probability), whereupon a small blue dot (0.15 dva) appeared on the white fixated circle, signalling that a saccade should be made to a yellow location within two seconds. The participant maintained fixation at the new location for 0.6 s, before the choices appeared for the next step. Steps proceeded in this manner until the goal was reached, after two, four, or six steps, whereupon the fixated circle turned blue for 1 s. At each step, if possible, the participant was to choose a yellow option that moved closer to the goal. In the simplest two-step routes, one of the two choice options for the first step was a move one step closer to the goal, and options for the second step included the goal itself (Figure 2a). In more demanding routes, some steps had two choice options that were equally attractive, either both towards or both away from the goal. Sometimes these longer routes involved a move into the outer perimeter of the maze.

For control problems, no goal location was shown at problem onset (Figure 2b, bottom). For 1.2 s, the participant simply maintained fixation on the central white dot. At each subsequent step, only one yellow circle was presented, cueing the move to be selected. Accordingly, the participant was to follow the route prescribed by this sequence of cued moves. Other details copied those for maze problems. The problem completed after 2, 4 or 6 moves, when the route had always led to one of the same four locations used as goals in maze problems, and when this location was reached, the fixated circle turned blue.

In fixation blocks, neither the goal location nor choice locations appeared, requiring the participant to maintain fixation on the central white circle for the entire problem, which was matched in duration to the mean duration of either two-, four-, or six-step problems, until the fixated circle turned blue.

In all block types, to maintain participant engagement, completing a problem was rewarded by a pleasant sound, and animated stars that travelled from the fixated location to the center of the grid. Every fourth problem had three stars, while other problems had one star, although this was of no significance. The movement of the stars encouraged gaze to return to the center, ready for the next problem, which started 0.5 s later. If any error was made (failing to saccade to a yellow circle in time, making a saccade before the go cue appeared, making a saccade to a red circle, or failing to choose the shortest available route to the goal), the current problem aborted immediately, with a central red cross and a disappointed sound (1 s), followed by the usual 0.5 s inter-problem interval. In the maze condition, three participants consistently failed to select the shortest available route to the goal (on >45% of problems, compared to <15% for other participants), so were excluded from all analyses.

Across problems of each condition, routes (defined by goal locations crossed with the location of choice options) were presented in pseudorandom order. On average, combining maze and control conditions, a run contained 14.1 two-step problems, 23.3 four-step problems and 18.8 6-step problems (with additionally 14.5 fixation problems).

Participants practised a few problems of each type outside of the scanner, using manual responses, until they and the experimenter were happy that they understood the task. In the scanner, the eye tracker was calibrated, and the participant practised responding using the button box and using saccades, before proceeding to the main experiment.

### Human: MRI acquisition

Scanning was performed on a 3 Tesla Siemens Prisma MRI scanner, using a 32-channel head coil. A T1-weighted structural volume was acquired using a 3D MPRAGE sequence (TR = 2.4 s, TE = 0.0022 s, flip angle = 8°, 0.8 mm isotropic voxels). A T2-weighted structural volume was acquired using a 3D SPACE sequence (TR = 3.2 s, TE = 0.56 s, flip angle = 120°, 0.8 mm isotropic voxels). During each of four task runs, T2*-weighted scans were acquired using a gradient echo 2D EPI sequence (TR = 0.80 s, TE = 0.037 s, flip angle = 52°, 2 mm isotropic voxels, 72 slices acquired with multiband factor 8). EPI runs were acquired with alternating phase-encoding directions (AP/PA). Two spin echo volumes, with similar phase reversal, were acquired between the second and third runs.

### Human: MRI analysis

Data processing was based on the HCP minimal pre-processing pipelines ((Glasser et al., 2013); version 4.7.0), called using automatic analysis software (Cusack et al., 2015). Per-subject processing steps are summarised briefly here. Structural volumes were undistorted using the spin-echo volumes, and rigidly aligned to MNI space. Subcortical structures and the cortical ribbon were segmented using the structural volumes, and cortical surface meshes were extracted, using FreeSurfer ((Fischl, 2012); version 6.0.0). Functional volumes were aligned to the structurals, undistorted using the spin-echo volumes, motion-corrected using FSL’s MCFLIRT (Jenkinson et al., 2002), and mapped to CIFTI “grayordinate” space, where they were smoothed with a 2 mm FWHM Gaussian kernel. Functional data were linearly detrended, and denoised using spatial ICA+FIX (Salimi-Khorshidi et al., 2014) on concatenated runs, with a FIX-threshold of 20. Cortical surfaces were registered to template space using multimodal surface matching (“MSMAll”; (Robinson et al., 2018)).

To model task-based activations, for each run, a first-level general linear model (GLM) was estimated using FSL’s FEAT (Woolrich et al., 2001). One regressor was defined for problems of each condition (whether correctly completed or not), as boxcars from initial fixation to the onset of feedback. These regressors were convolved with the canonical haemodynamic response function (HRF) and its temporal derivative. A temporal high-pass filter with a cut-off period of 200 s was applied to the data and the model. Activation due to goal-based decisions was estimated from the parameter estimates for the maze condition’s HRF-convolved regressor contrasted against the control condition. Statistical significance was tested across participants, using two-tailed, one-sample t-tests. This was initially performed at every cortical vertex, then repeated for each cortical region in the HCP’s multimodal parcellation (Glasser et al., 2016), after averaging across vertices and hemispheres for each region. In both analyses, multiple comparisons were accounted for by controlling the false discovery rate (FDR = 0.05) (Genovese et al., 2002). Results were further summarised at a network level according to the networks defined by (Ji et al., 2019) (excluding parcels whose network assignment differed across hemispheres).

A supplementary analysis (Supplementary Figure 1) contrasted just two-step maze with two-step control problems, with four-and six-step problems removed as regressors of no interest.

### Monkey: Participants

Subjects were two male rhesus monkeys (Macaca mulatta), each weighing 14 kg. Experiments were performed in accordance with the Animals (Scientific Procedures) Act 1986 of the UK; all procedures were licensed by a Home Office Project License obtained after review by Oxford University’s Animal Care and Ethical Review committee, and were in compliance with the guidelines of the European Community for the care and use of laboratory animals (EUVD, European Union directive 86/609/EEC).

For each animal, an MRI-compatible head holder (Rogue Research) was sterotaxically implanted on the skull. All surgical procedures were aseptic and carried out under general anesthesia. At the end of the experiments, animals were deeply anesthetized with barbiturate and then perfused through the heart with heparinized saline followed by 10% formaldehyde in saline.

### Monkey: Maze task

For monkeys, the task was a simplified version of the human maze. Problems were presented on a grid of lines (Figure 2c), corresponding just to the inner 3 x 3 locations of the human maze. In the maze condition, all routes were two-step, with one of the two choice options at step 1 moving closer to the goal, and the goal then reached at step 2. In the control condition, again, moves were forced by just a single available option at each step.

Timings for the maze condition are illustrated in Figure 2d. At the start of each problem, subjects were given a green fixation point in the middle of the grid and had to fixate on this point within 2 s. Once the fixation was initiated, a white diamond (cursor) appeared in the location of the fixation. After 0.2 s a blue circle (final target) appeared at the final goal location of the current problem and remained on the screen for 0.5 s while the monkeys kept fixating in the center (except for a single session, when the target was shown for 0.4 s).

The final goal was only shown at this point and then disappeared, so the monkeys needed to hold the location in mind throughout the remainder of the problem. After a 0.3 s delay, the 4 choice options of the first step were presented. 2 green dots and 2 red dots appeared simultaneously at the 4 nodes next to the central one (where the monkeys were still fixating). The red dots represented nodes that were blocked, and only one of the greens was a correct choice, being closer to the final target location. The go signal to make a saccade was the disappearance of the green fixation point, occurring after a delay randomly either 0.3 or 0.9 s. After the go signal, the monkey had 0.5 s to initiate a saccade to the correct choice location and 0.05 s to complete the saccade. Once a saccade to the new location was made, if the choice was correct, all the other old stimuli (initial fixation point, cursor, other choices) disappeared while the monkeys had to hold fixation on the new location.

The second step of the problem had a similar structure: the cursor appeared when the second fixation started, the second set of 3 choices (2 green 1 red) appeared after 0.3 s and were presented for the same random delay as before, followed by the go signal. Using the same timing as in step 1, the monkey needed to make a saccade towards the correct green location. At this point, one of the choice options lay on the actual final goal location, allowing the monkeys to complete the problem. If this happened, when the eye landed, the cursor appeared at the fixated target location, joined after 0.05 s by brief (0.05 s) reappearance of the blue target itself. A smoothie reward was then delivered (delivery duration: ∼0.05 s), the monkey could break fixation, and the problem ended with the disappearance of all the stimuli (except the grid) from the screen and an inter-problem interval of 1 s. Errors at any point during the problem (incorrect location selections, timing errors, fixation breaks) led to an immediate error cue (large red circle) shown in the center of the screen, and termination of the problem without reward.

Events for the control condition were similar, except that no target was shown at the start of the problem. Again, each problem had two steps, each with only a single choice location, and reward was delivered after the second correctly executed step. In an additional fixation condition (not analysed here), each problem began with a green fixation point in the middle of the grid. Fixation was required within 2 s, and once it was achieved, the fixation point was surrounded by a white diamond. Maintained fixation was required for at least 2.05 s, after which a blue circle appeared at fixation for 0.05 s, and reward was delivered.

As for the human, the experiment used a block design. For as long as the monkey kept working, blocks of the maze condition alternated with blocks of a non-maze condition. Each non-maze block was randomly selected to be either control or fixation, after which the next block was again maze. The target length of each block was 30 s. When a problem was completed, the block terminated if >24 s had already elapsed; otherwise a new problem began and the block continued until this problem was finished.

The task was programmed in Psychopy Release 2021.2.3 (https://www.psychopy.org/). Stimuli were presented on a 23 inch LCD screen (https://www.crsltd.com/tools-for-functional-imaging/mr-safe-displays/boldscreen-23-lcd-for-fmri/), positioned 80 cm from the monkey’s eye. The grid measured 15 x 15 dva. The fixation point and choice cues were filled circles measuring 1.15 dva. The cursor, blue target, and error cue measured 2.3 dva. Eye movement data were sampled at 1000 Hz using an MRI-compatible tracking system (EyeLink 1000 Plus, SR Research; https://www.sr-research.com/). At each step, fixation was required within a 7 x 7 dva window centered on the appropriate grid position. Reward (fruit smoothie) was delivered through a Peristaltic Electric Operated Positive Displacement Pump (Verderflex).

Prior to data acquisition, each monkey acquired the task in a series of stages. They were first trained on the basic oculomotor components of the task: holding central fixation, making saccades to targets, maintaining fixation at different locations on the screen, and making delayed saccades to a target. They then progressed to a one-step version of the final task and, after reaching sufficient accuracy, to the full two-step version. In parallel with the final phases of the task training, the animals were gradually acclimatised to working in the sphinx position and to the scanner environment, which required a separate and extended habituation period.

### Monkey: MRI acquisition

Data were collected over 16 sessions in monkey A, and 18 in monkey B (5 sessions from monkey B were excluded based on poor behavioral performance). Animals were head-fixed in a sphinx position in an MRI-compatible chair. fMRI data were collected using a whole-body 3T MRI scanner with a full-sized horizontal bore and a custom-made four-channel phased-array local receive coil in conjunction with a radial transmit coil (H. Kolster; Windmiller Kolster Scientific, Fresno, CA, USA). Whole-brain blood-oxygen-level-dependent (BOLD) fMRI data were acquired using a gradient echo T2* echo planar imaging (EPI) sequence with 1.5-mm3 isotropic voxel size resolution, 36 interleaved slices with no gap, repetition time (TR) = 2.28 s, echo time (TE) = 30 ms, and flip angle = 90°. Proton density– weighted images using a gradient-refocused echo (GRE) sequence (TR = 10 ms, TE = 2.52 ms, and flip angle = 25°) were acquired as references for body motion artefact correction. T1-weighted magnetization-prepared rapid gradient echo images (0.5-mm^3^ isotropic voxel size resolution, TR = 2.5 ms, TE = 4.01 ms, inversion pulse time = 1.1 s, and flip angle = 8°) were acquired in separate anesthetized scanning sessions. In these sessions, monkeys were scanned in sphinx position in an MRI-compatible stereotactic frame (Crist Instrument). Anesthesia was induced via intramuscular injection of ketamine (10 mg/kg), xylazine (0.125 to 0.25 mg/kg), and midazolam (0.1 mg/kg).

### Monkey: Preprocessing and reconstruction pipeline

Magnetic resonance images were preprocessed using tools from the FMRIB Software Library (FSL) and the Magnetic Resonance Comparative Anatomy Toolbox (MrCat; www.neuroecologylab.org). The GRE image was used to carry out the T2* EPI image reconstruction by an offline-SENSE reconstruction method (Dr. Kolster, Windmiller Kolster Scientific, Fresno, CA, USA).

### Monkey: MRI analysis

Data were analysed in Matlab (The Mathworks Inc.), using automatic analysis software (Cusack et al., 2015) to call SPM functions (Statistical Parametric Mapping; www.fil.ion.ucl.ac.uk/spm). The T1-weighted structural scan was spatially normalized to the 112RM-SL template (McLaren et al., 2009) using nonlinear warping, and segmented by tissue type (grey matter, white matter, CSF). The first ten EPI volumes of each session were discarded, and then the remaining volumes were rigidly realigned to each other, corrected for slice timing, and rigidly co-registered to the structural volume. Although the head was fixed, limb movements caused changes in the magnetic field, resulting in momentary spatial distortion and extreme intensity fluctuations. Time points affected by such artefacts were defined as those with an apparent displacement of the brain by > 1 mm, an apparent rotation of > 0.5 degrees around any axis, or an extreme global intensity difference from the previous scan. (Global intensity differences were calculated as the mean across voxels of the squared difference between one volume and the next; this time-course was smoothed with Matlab’s robust Lowess method, with a span of 10% of the data length; outliers were defined as exceeding four standard deviations of the lower 90% of the distribution of residuals from the smoothed time-course.) EPI volumes were finally warped to template space, using the parameters derived from the structural, resliced to 1 mm isotropic resolution, and spatially smoothed using a 4 mm FWHM Gaussian kernel.

Because animals often broke fixation when not working well, the analysis included just problems that did not terminate with a timing error. For each session, the first-level GLM included one regressor to model problems of the maze condition, another for the control condition, and similar, but unanalysed, regressors to model excluded problems. These regressors were constructed from boxcars spanning initial fixation to the onset of feedback for each problem, convolved with SPM’s canonical HRF. A separate regressor modelled moments when any juice rewards were delivered, timed for 1 s from the recorded triggers, and convolved with the HRF. To account for variance due to limb movements, the model also included extensive unconvolved covariates of no interest. These included the six rigid-body realignment parameters, their squares, and temporal derivatives of both, as well as the mean signal from the CSF, and individual regressors to model any scan identified as a global outlier (as defined above). A temporal high-pass filter with SPM’s default cut-off period of 128 s was applied to the data and the model. Activation due to goal-based decisions was estimated from the parameter estimates for the maze condition’s HRF-convolved regressor contrasted against the control condition. Statistical significance, per voxel, was tested across sessions x hemispheres, using two-tailed t-tests, based on a repeated-measures ANOVA with separate covariates to model each hemisphere of each animal, and a pooled error term that assumed similar error covariance across voxels (Henson, 2015). Multiple comparisons were accounted for by controlling the false discovery rate (FDR = 0.05).

To warp macaque surface-based activations to the human cortical surface, we used the transformation published by (Xu et al., 2020). This required two steps: the activations were first warped from the F99 surface to the Yerkes19 surface, using the transformation in RheMAP-Surf (Zhou and Xu, 2025), and then from the Yerkes19 surface to the human surface (Xu et al., 2020). Both steps were performed using Connectome Workbench version 1.5.0.

## Acknowledgments

This work was supported by Wellcome Trust Grant 101092/Z/13/Z, Wellcome Trust Early Career Award 305264/Z/23/Z, Medical Research Council UK Program MC_UU_00030/7, and Carlsberg Foundation Grant CF22-1111. We thank Simon Baumann and Stuart Mason for assistance with monkey training, and Makoto Kusunoki for surgery. For the purpose of open access, the author has applied a Creative Commons Attribution (CC BY) licence to any Author Accepted Manuscript version arising from this submission.

## Supplementary materials

**Supplementary Figure 1.**
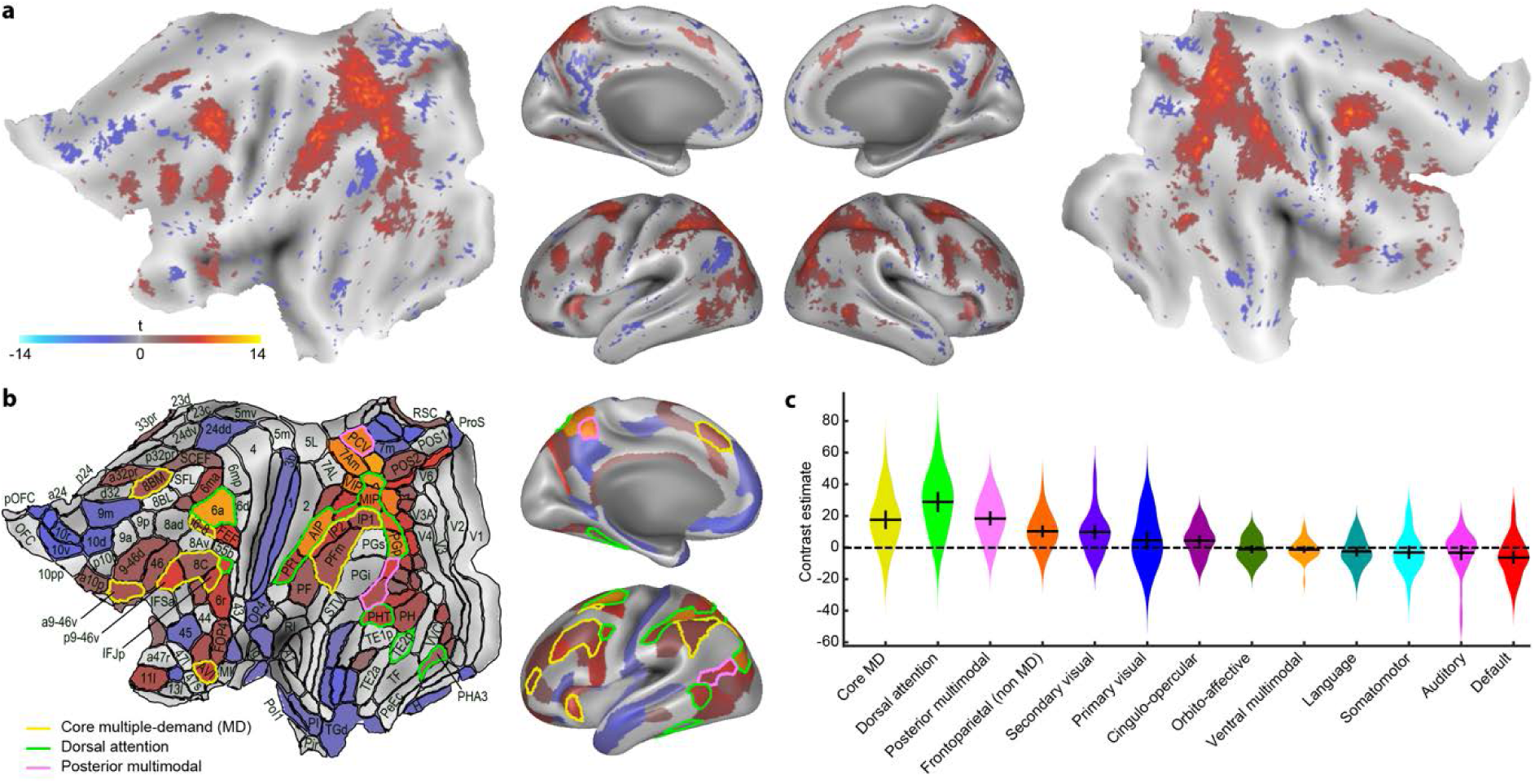
Contrast of maze minus control: Human data, two-step problems only. Details as Figure 4.

**Supplementary Figure 2.**
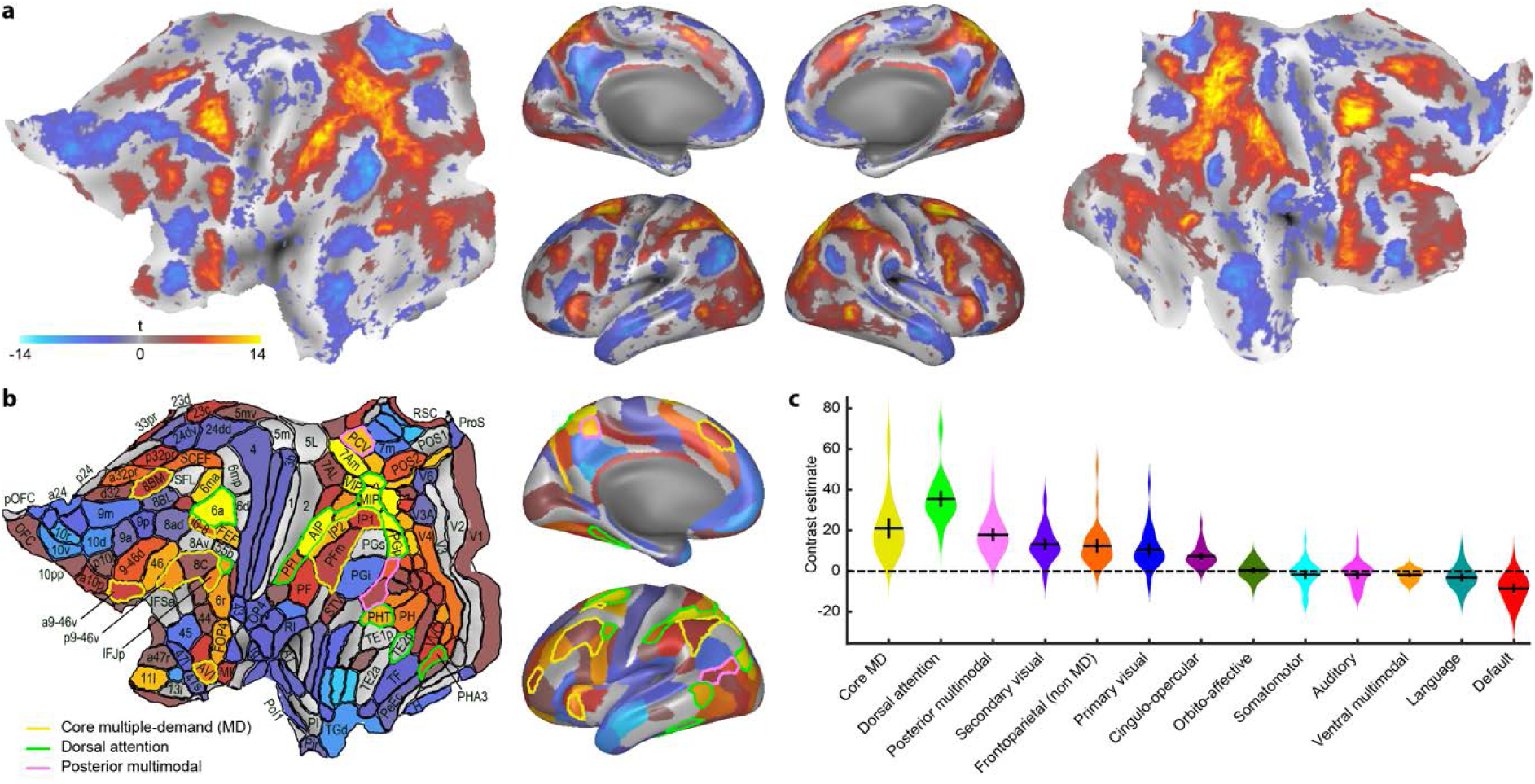
Contrast of maze minus control: Human data, manual-response task. Details as Figure 4.

**Supplementary Figure 3.**
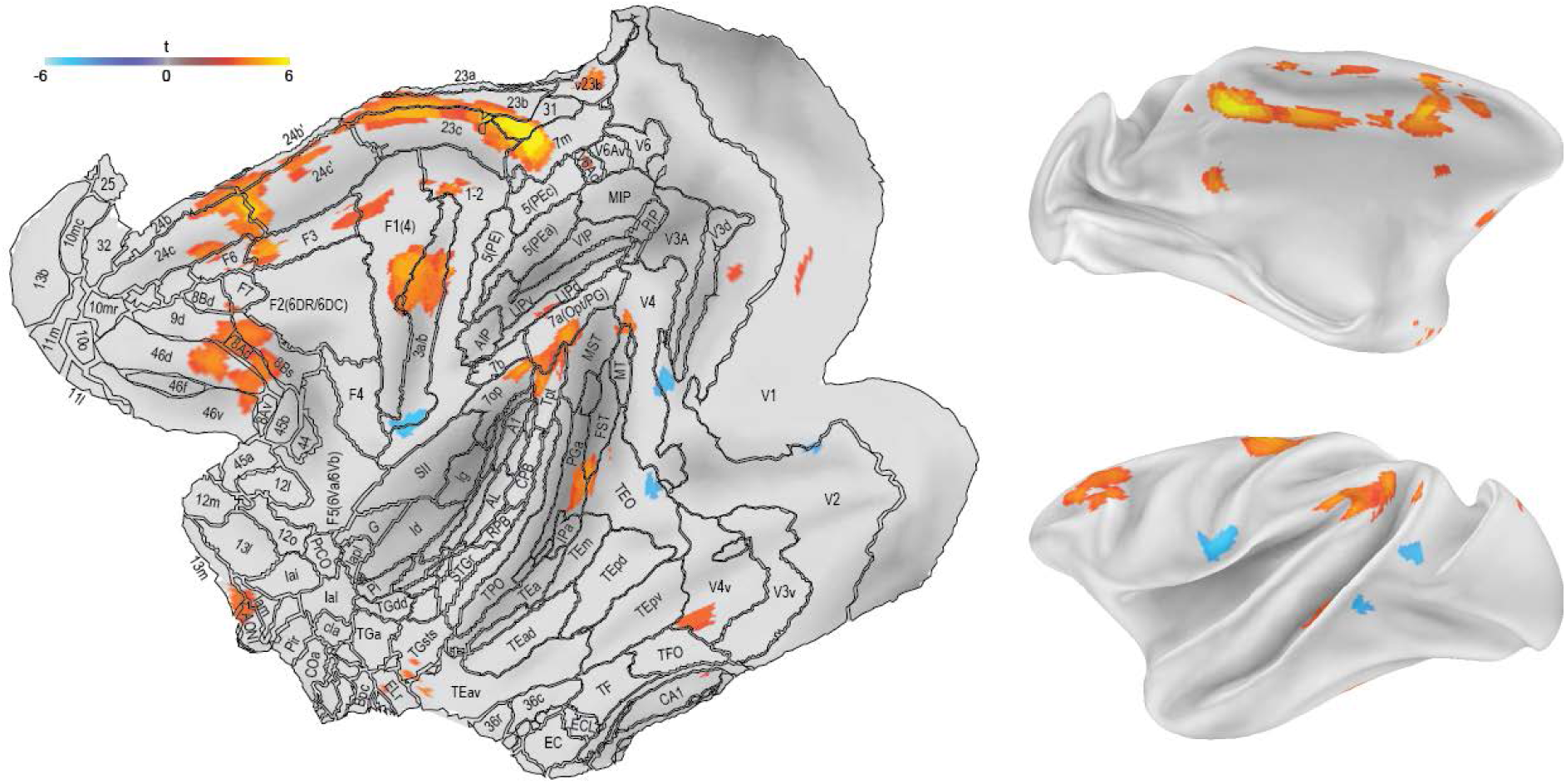
Contrast of maze minus control: Macaque data, correctly completed problems only. Details as Figure 5.

**Supplementary Figure 4.**
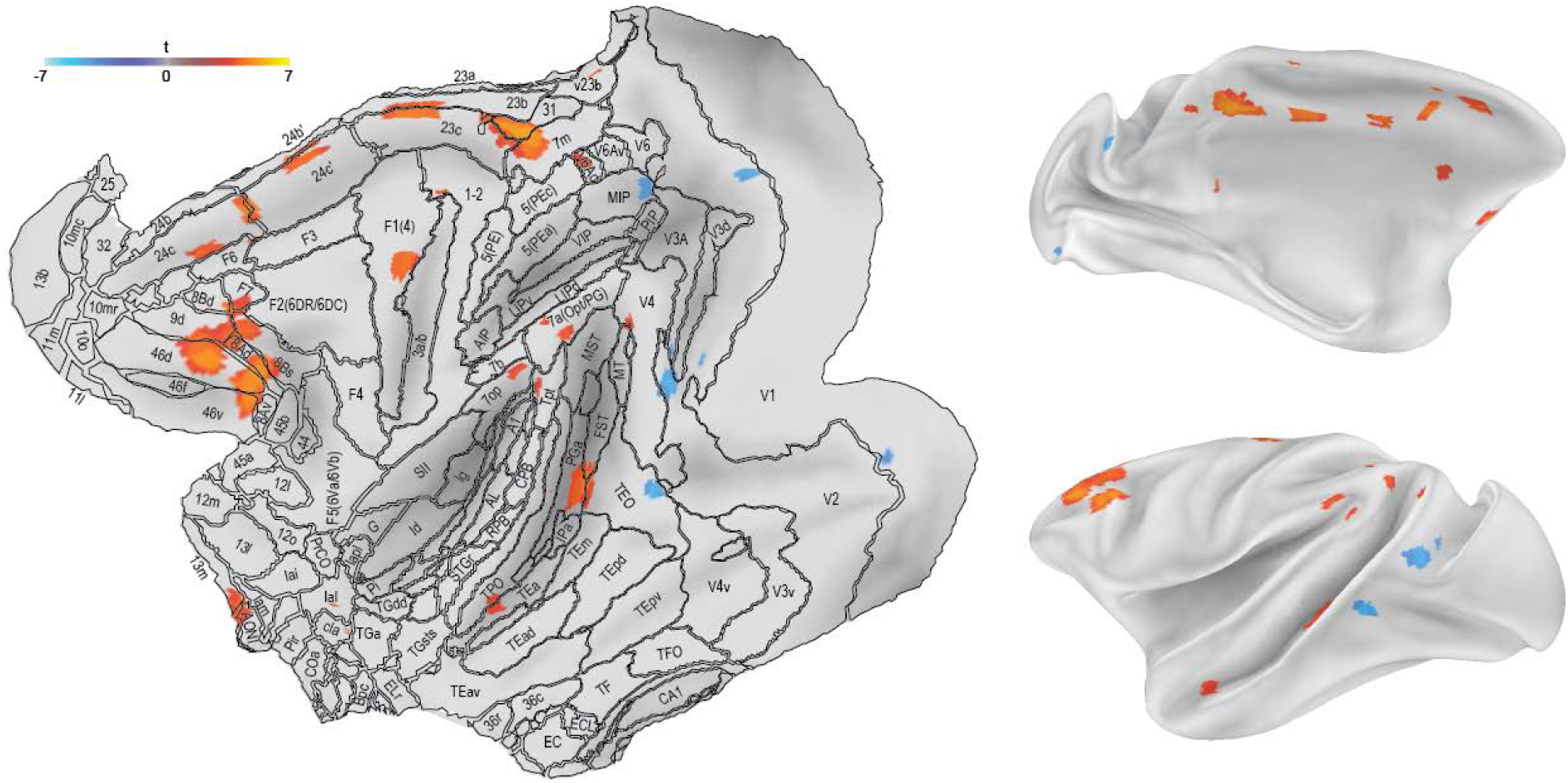
Contrast of maze minus control: Macaque data, 3 mm FWHM smoothing kernel. Details as Figure 5.

**Supplementary Figure 5.**
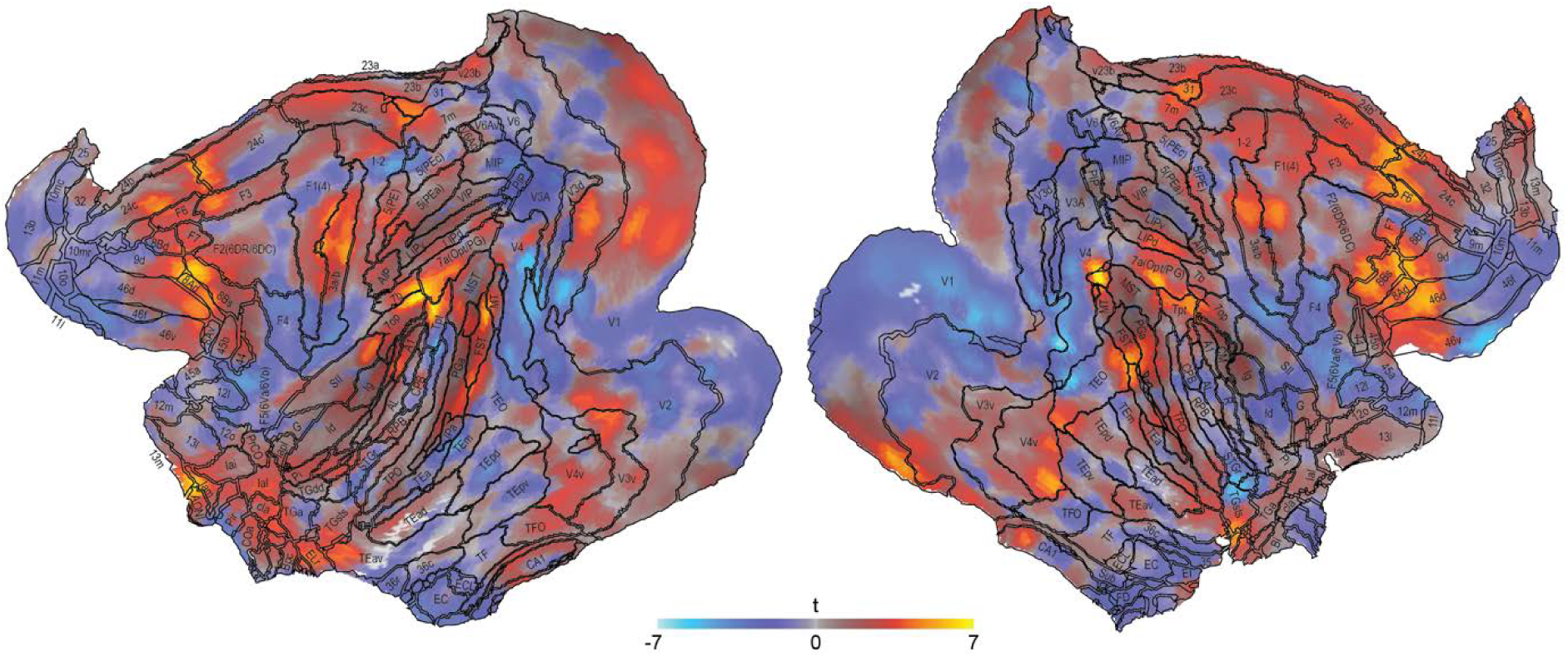
Contrast of maze minus control: Macaque data. Separate data for left and right hemispheres, averaged over animals.

**Supplementary Figure 6.**
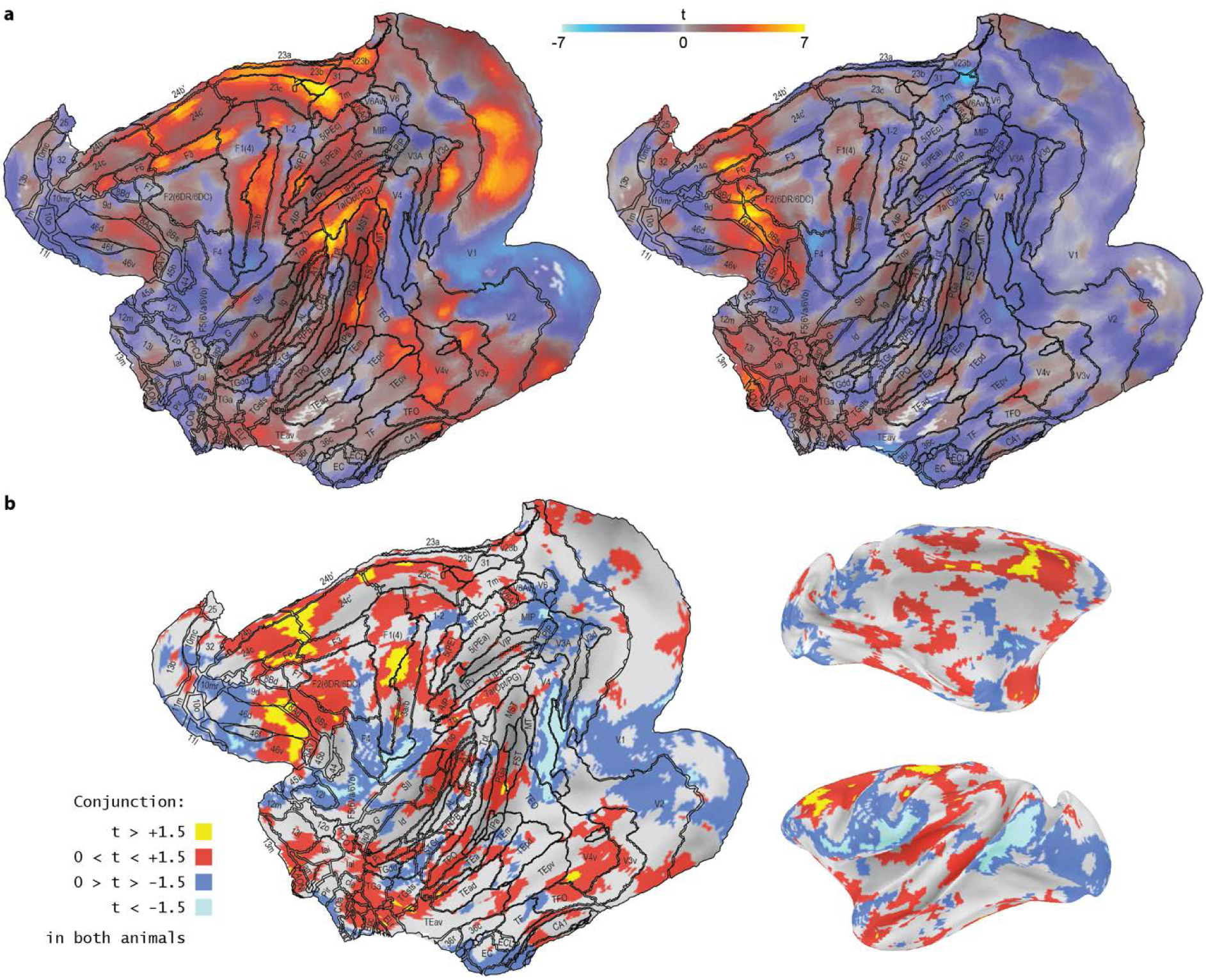
Contrast of maze minus control: Macaque data. (a) Separate data for monkeys A and B, averaged over hemispheres. (b) Conjunction map of data in (a).

