## Supplementary Materials for "Cognitive control networks in human and macaque"

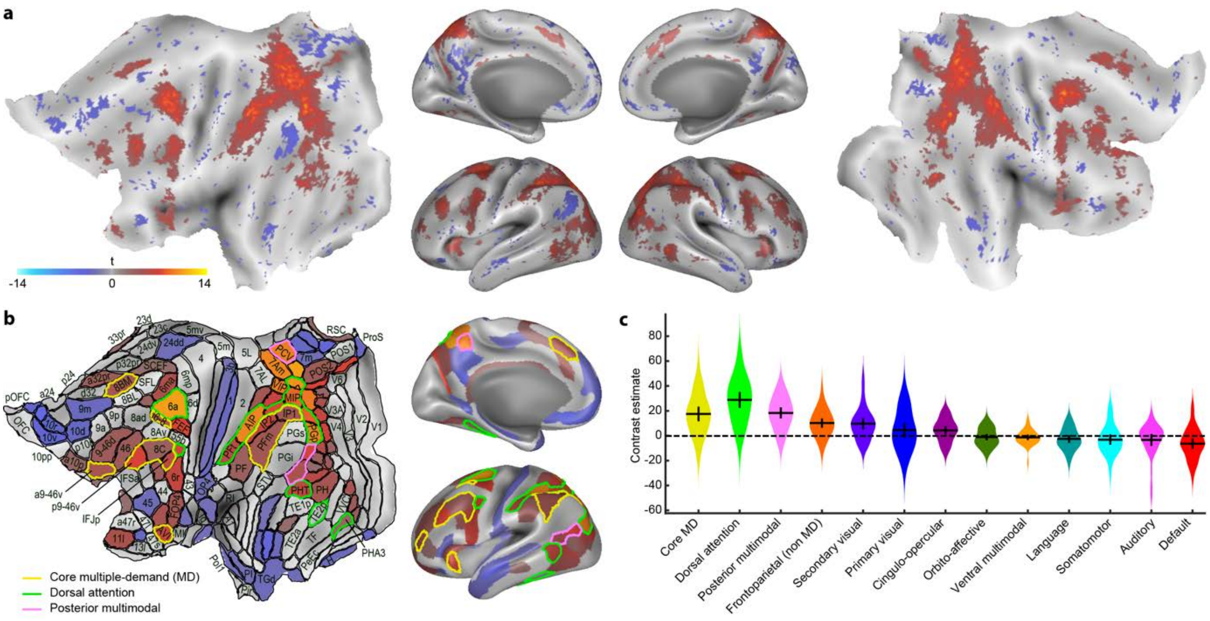


Supplementary Figure 1. Contrast of maze minus control: Human data, two-step problems only. Details as Figure 4.

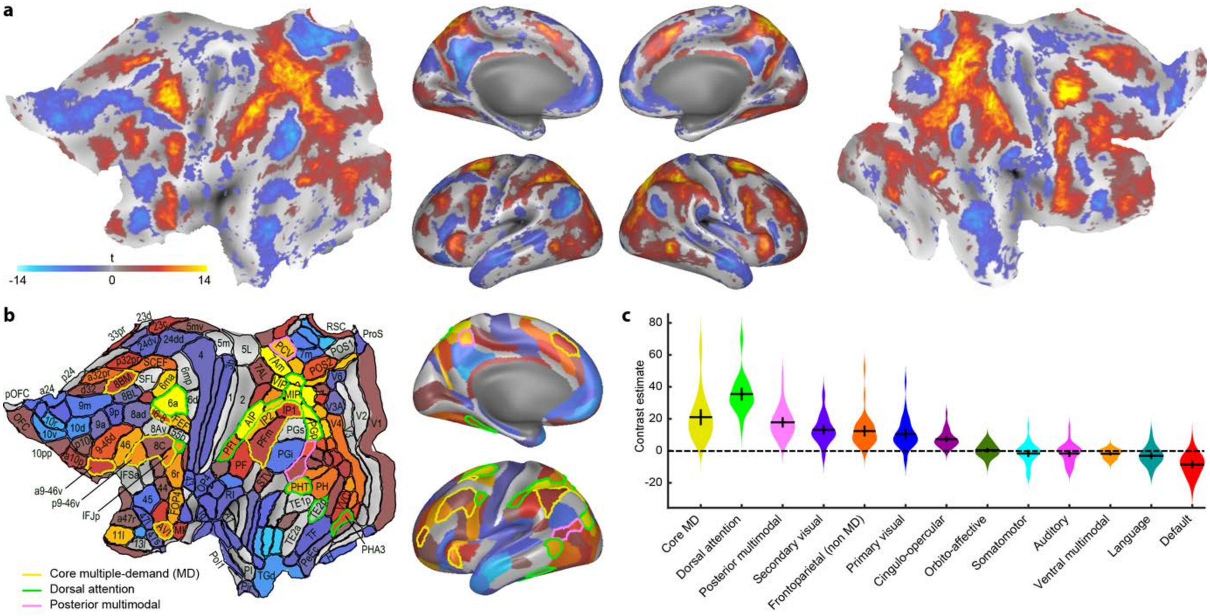


Supplementary Figure 2. Contrast of maze minus control: Human data, manual-response task. Details as Figure 4.

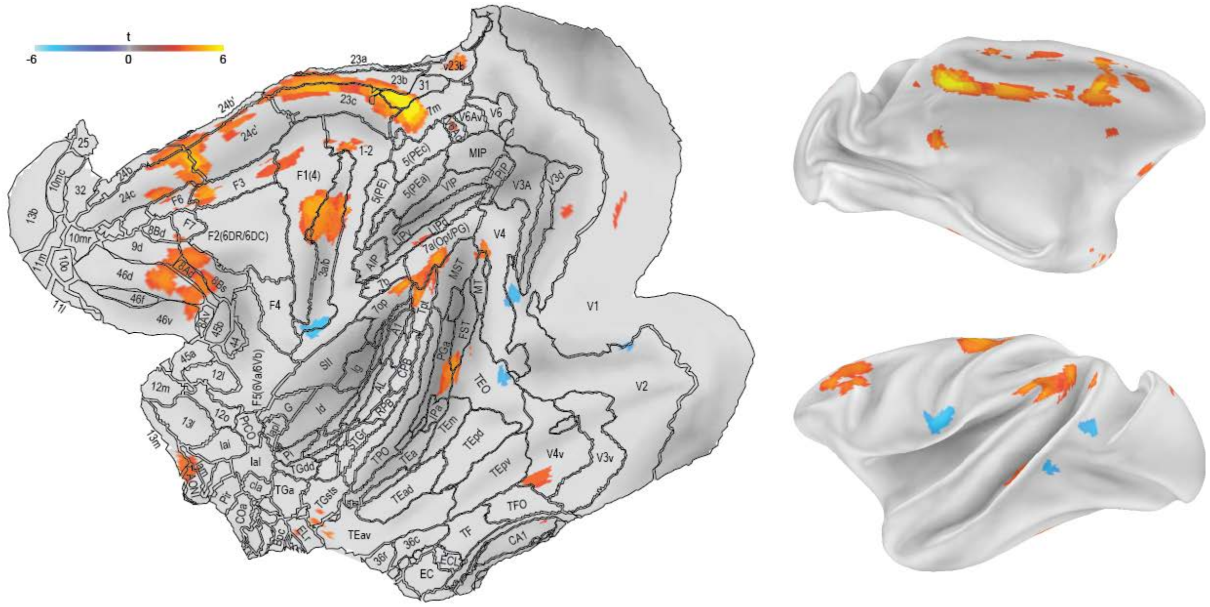


<u>Supplementary Figure 3</u>. Contrast of maze minus control: Macaque data, correctly completed problems only. Details as Figure 5.

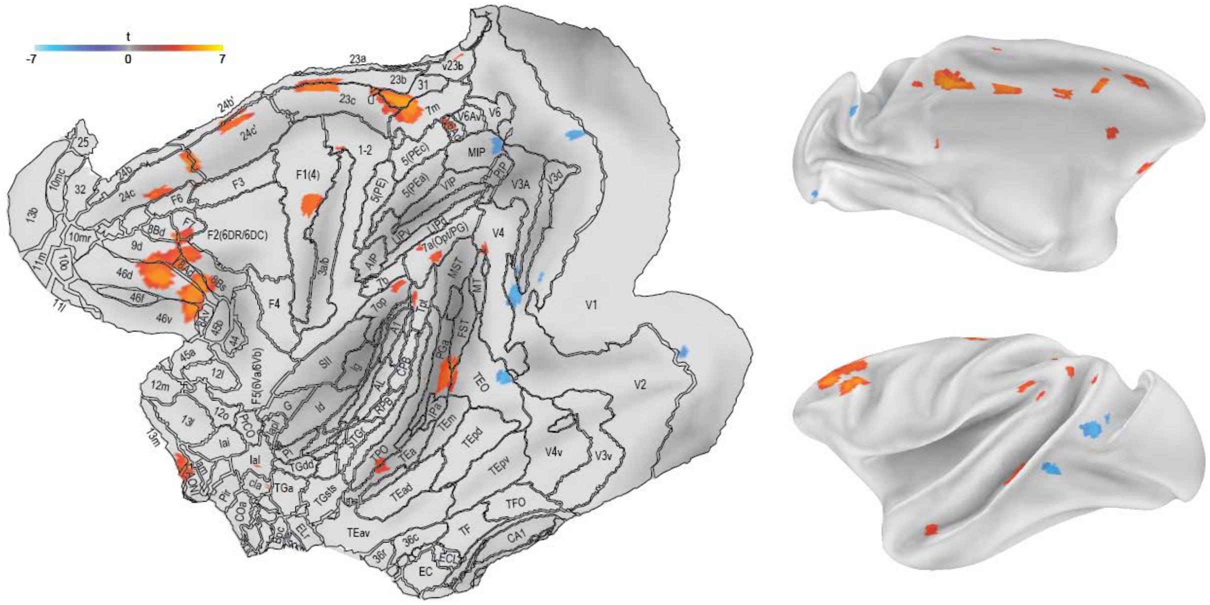


<u>Supplementary Figure 4</u>. Contrast of maze minus control: Macaque data, 3 mm FWHM smoothing kernel. Details as Figure 5.

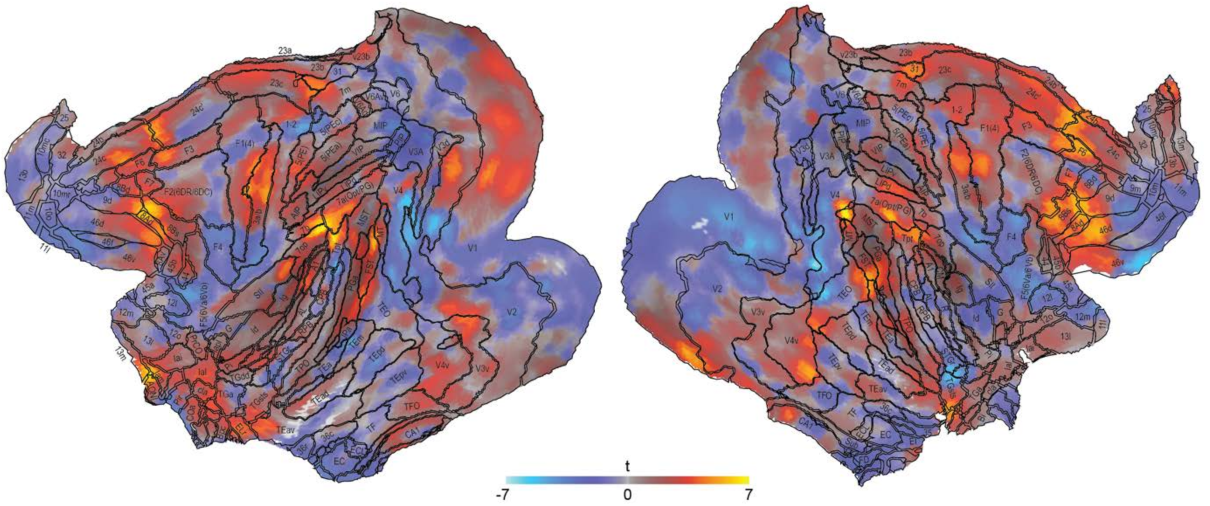


Supplementary Figure 5. Contrast of maze minus control: Macaque data. Separate data for left and right hemispheres, averaged over animals.

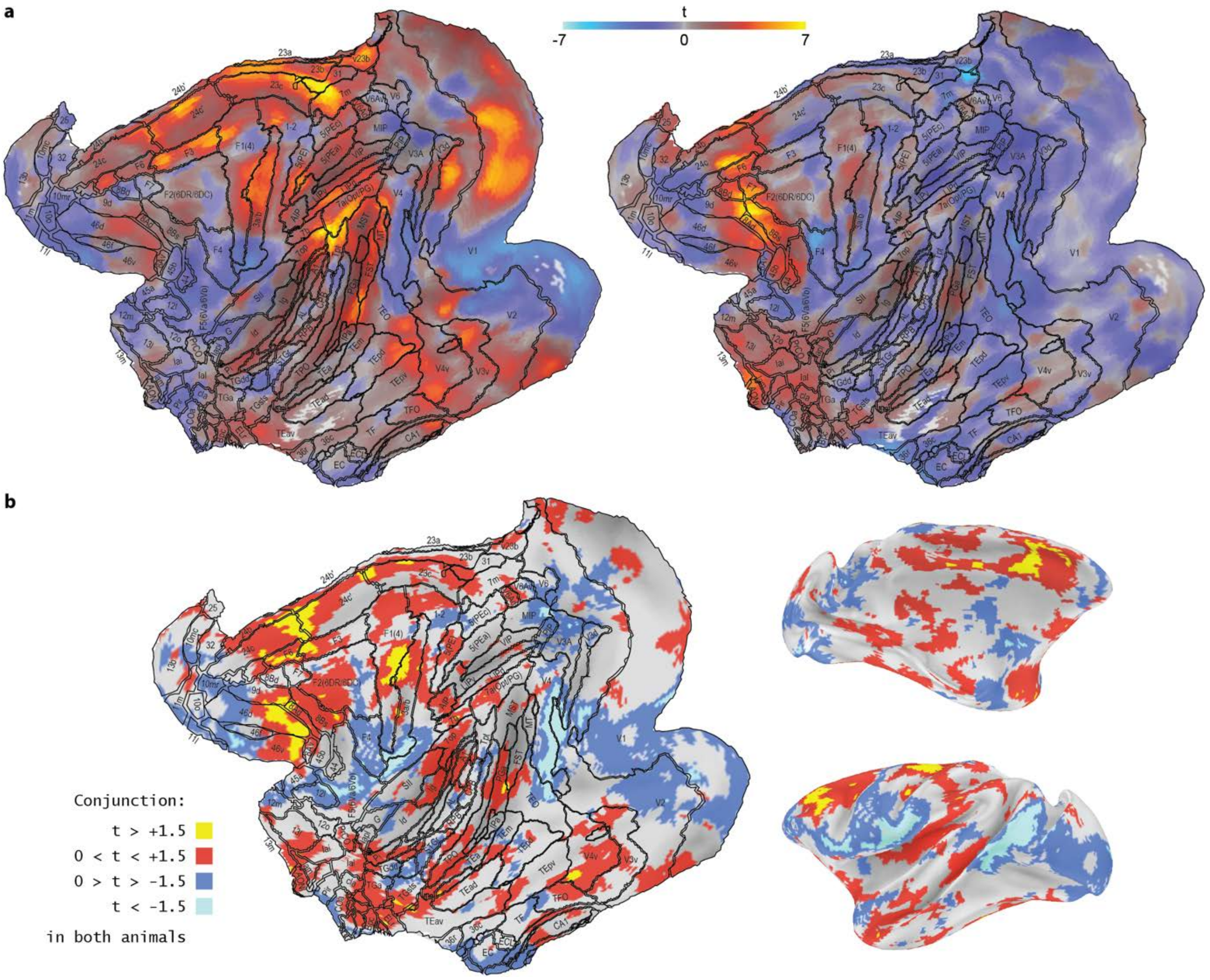


Supplementary Figure 6. Contrast of maze minus control: Macaque data. (a) Separate data for monkeys A and B, averaged over hemispheres. (b) Conjunction map of data in (a).
